# Signatures of processing complexity during global cognitive states in ventromedial prefrontal cortex

**DOI:** 10.1101/2020.10.08.331579

**Authors:** Priyanka S. Mehta, Seng Bum Michael Yoo, Benjamin Y. Hayden

## Abstract

Behavioral neuroscience almost exclusively studies behavior during tasks and ignores the unstructured inter-trial interval (ITI). However, it is unlikely that the ITI is simply an idling or paused mode; instead, it is a likely time for globally focused cognition, in which attention is disengaged from the task at hand and oriented more broadly. To gain insight into the computational underpinnings of globally focused cognition, we recorded from neurons in a core decision-making region, area 14 of ventromedial prefrontal cortex (vmPFC), as macaques performed a foraging search task with long inter-trial intervals (ITIs). We find that during the ITI, ensemble firing is associated with increased discriminability of a key mnemonic variable, recent reward rate, which in turn predicts upcoming search strategy. ITI activity is also associated with increased ensemble dimensionality and faster subspace reorganization, presumed markers of processing complexity. These results demonstrate the flexible nature of mnemonic processing and support the idea that the brain makes use of ostensible downtime to engage in complex processing.

## INTRODUCTION

Our minds can readily switch between states that are stimulus-focused and oriented narrowly on the task at hand (***local states***) and stimulus-independent states that are relatively undirected and more attuned to the world around us (***global states***, Raichle, 2010; Raichle, 2015; Mason et al., 2007; Smallwood et al., 2013). During local states, many of the task-irrelevant things on our mind fall away so that we can selectively process the most important and urgent information. When our thoughts move away from the task, our minds then fill with a variety of stimulus-independent thoughts, including recollections of the recent past, as well as background and off-task cognition (Smallwood et al., 2013; Baird et al., 2013; Konishi et al., 2015).

Globally oriented cognition is poorly understood. We know a lot more about what goes on in the brain during brief periods of focused performance than during the often much longer times between. However, far from being a mental equivalent of doing nothing, globally oriented cognition appears to be rich and complex, and too extend both backwards and forwards in time (Northoff et. al, 2010, Mazoyer et al., 2001; Zou et al., 2013; Cole et al., 2016). Indeed, even if global cognition is not directly related to the specific details of the most recent trial, it is associated with several functions, such as reactivating past sequences of action, guiding navigation in future goal-direction movement, and maintaining cognitive control and working memory (Olafsdottir et al., 2018; Raichle, 2010; Jilka et al., 2014;). One theme that unites these functions is that they have at least some mnemonic relevance. Specifically, they involve reactivation of memories of the recent and distant past.

Global cognitive functions are particularly associated with a discrete network of brain regions known as the default mode network (DMN). Hemodynamic activity in DMN regions is enhanced during delays between trials and in the absence of specific task instructions (Raichle et al., 2015; Behrens et al., 2018; Hayden et al., 2009b) Activity in the DMN appears to reflect cognitive processes that are antagonistic to task focus. For example, DMN activity predicts sporadic lapses in attention (Weissman et al., 2006), failures to encode memories (Daselaar et al., 2004), and failures to perceive near-threshold somatosensory stimuli (Boly et al., 2007). One region of the DMN, the medial prefrontal cortex, is associated with an increased BOLD signal when subjects are tasked with processing information from previous trials (Konishi et al. 2015) including memory consolidation during stimulus-independent thought (Smallwood et al., 2013; Baird et al., 2013). These observations support the idea that activation of mnemonic processes naturally competes with effective task performance.

We wished to gain insight into the neural processes occurring in global cognition. We examined ensemble firing rates in one DMN structure, the ventromedial prefrontal cortex (vmPFC), as macaques performed a computerized foraging task. The task, like most laboratory tasks, is naturally divided into a task period and an inter-trial interval (ITI). Our analyses therefore centered on comparing neural activity during the task to activity during the ITI. We used a computerized foraging task that dissociates behavioral modes in macaques. On each trial (the presumed local state), the subject must search for several seconds between options that offer obscured riskless but variable rewards. During the 4-second ITI subjects presumably enter into a globally-oriented cognitive state. To gain insight into the role of vmPFC in global cognition, we focused on a behavior known as *adjustment*. Adjustment is a form of rapid learning and may reflect a strategic change motivated by an underlying belief that task statistics have changed (Hayden et al., 2008; Hayden et al., 2009a; Hayden et al., 2011; Blanchard et al., 2015). We hypothesized that the ITI may be a critical period for performing adjustment-related computations.

We find three striking differences between these two task phases. First, information about the rate of recent reward intake (over the past 10 trials) is more readily distinguished in the ITI, and that this information closely predicts subsequent adjustment. Second, neural ensemble dimensionality is substantially greater during the ITI. Third, the *change rate* of the subspace reorganization is significantly enhanced during the ITI. Together, these results highlight the inherent richness of neuronal processing during the ITI, and during global cognition more generally, and, in particular, suggest that this processing privileges mnemonically relevant information.

## RESULTS

Macaques performed a foraging search task in which they inspected a series of hidden offers and selected one of them to obtain a liquid reward (**Figure 1A**, Mehta et al., 2019). The reward was deterministic but its value was randomized by offer and not predictable before inspection. Search in this task is self-terminating, so the task has features in common with the secretary problem and other stopping problems (Ferguson, 1989). Detailed behavior in the task is summarized in our previous paper, is not relevant to our hypotheses, and is not repeated here (Mehta et al., 2019).

**Figure 1.**
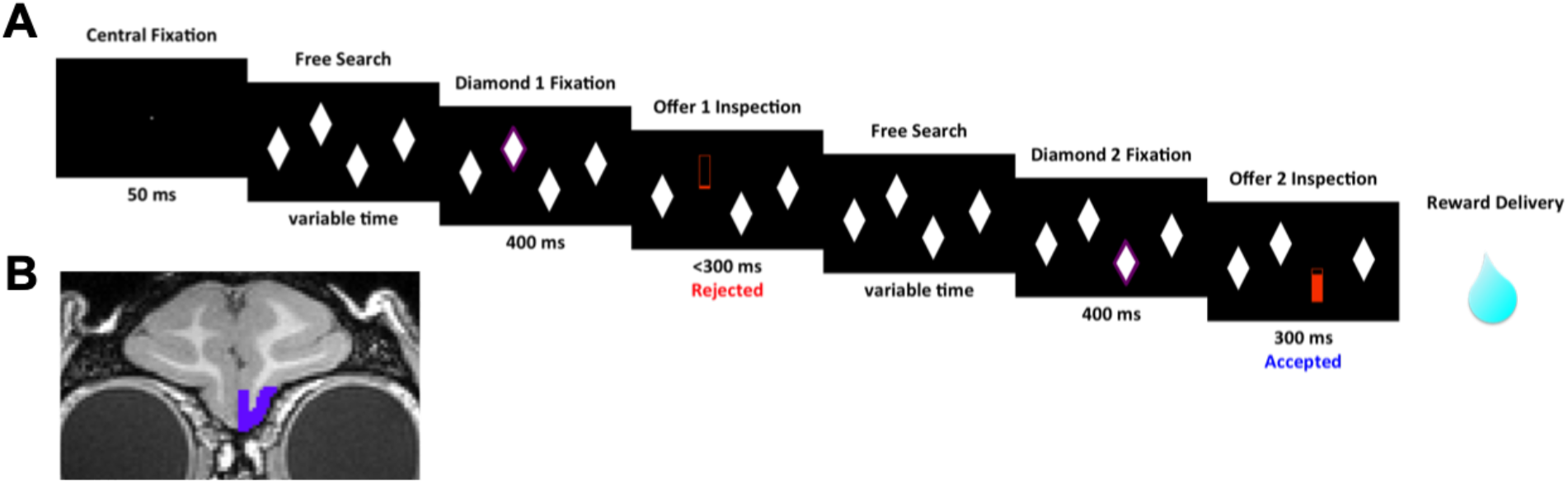
Our foraging task. **(A)** A typical trial. Subject searches through multiple offers and accepts one, ending the trial. **(B)** Area from which all recordings were obtained: macaque area 14, analogue of human ventromedial prefrontal cortex.

While our subjects performed the task, we recorded responses of 122 single neurons in area 14 of vmPFC (**Figure 1B**). We found that individual neurons encode several foraging-related variables during the trial, including the value of the current offer being inspected, the value of the most recent offer inspected, the outcome of the previous trial, the local reward rate, and the location of the attended offer (Mehta et al., 2019). While our previous study focused on trial-related activity, here we focus primarily on comparing activity between the trial and the inter-trial interval (ITI). The key feature of our search task that enables this analysis is the unusually long ITI (4 seconds), which allowed us to study responses temporally dissociated (>1 second) from both the past trial and the upcoming one, thus avoiding both sustained / hysteretic and anticipatory effects.

We focused our analyses on two 500 ms epochs. The *ITI epoch* is centered on the midpoint of the ITI - exactly two seconds after it begins and two seconds before it ends. This means our window of analysis spans 250 ms before this point, and 250 ms after. Our *trial epoch* is centered on the midpoint of the 1-second time period after the reveal of the first offer of each trial. This window of analysis spans from 250 ms after the offer reveal to 750 ms after the offer reveal.

### Mnemonic information is more decodable during the ITI than during the task

Our central hypothesis is that firing patterns during the inter-trial interval are neither functionally irrelevant nor otiose, but instead have a rich functional repertoire that differs from that of the task period. Based on existing literature, we hypothesized that functionally meaningful activity during the ITI would be associated with an expressly mnemonic role. In this task, rapid learning takes place on the basis of recent outcomes (Mehta et al., 2019). This adjustment may reflect a strategic response to evidence that environmental richness has changed (cf. Stephens & Krebs, 1986), or it may simply reflect a win-stay/lose-shift-like behavior. In either case, in this task *recent reward rate* (specifically, the average reward obtained over the past ten trials), influences the subject’s behavioral threshold on the current trial (□ = 0.2313, p< 0.001, linear regression of combined data across subjects between past 10 trials excluding most recent one against observed behavioral threshold). Note that this effect is not simply due to single trial adjustment (cf. Hayden et al., 2008; Sugrue et al., 2004). We know this because our analysis excludes the most recent trial. Thus, all the events that drive the response come from 10- 70 seconds before the choice is expressed.

Recent reward rate is robustly encoded by responses of vmPFC neurons. Indeed, during the 500 ms trial epoch, 22.13% of neurons in vmPFC encoded the average reward rate on the past ten trials (18.57% and 26.92% in subjects J and T, respectively). (As above, this and the next analysis exclude the most recent trial to eliminate bias by possible behaviorally hysteretic effects). A similar proportion of neurons (23.77%) encode the same information during the 500 ms ITI epoch (J: 28.57%; T:17.31%). All four of these proportions are substantially greater than chance (binomial test, p<0.0001 in all cases).

We next hypothesized that, if ITI was associated with mnemonic function, then the recent reward rate would be more *informationally accessible* during the ITI than during the task period. To test this hypothesis, we first devised a measure of informational accessibility based closely on a technique previously devised by Cohen and Maunsell (2010) and extended by Keemink and Machens (2019). We used a dimensionality reduction approach; specifically, one that uses Principal Component Analysis (PCA). We first projected our condition-averaged peri-stimulus time histograms (which represent time binned vectors of firing) into a PCA space. That is, we averaged all the peri-stimulus time histograms in an epoch across each condition (in this case, high and low reward rate).

In this way, we mathematically reduced our set of firing rates down to a set of *principal components* that efficiently describe the population activity as it varies with reward rate. We focused on the three principal components that describe the largest amount of neural activity for each group of trials (high-reward-rate trials and low-reward-rate trials). This allowed us to graphically represent the population activity: each principal component became an axis, with each point on the resulting three-dimensional plot representing a projection of the neural population (**Figure 2A**). We compared the two sets of points - the neural population during low-reward-rate trials versus the neural population during high-reward-rate trials.

**Figure 2.**
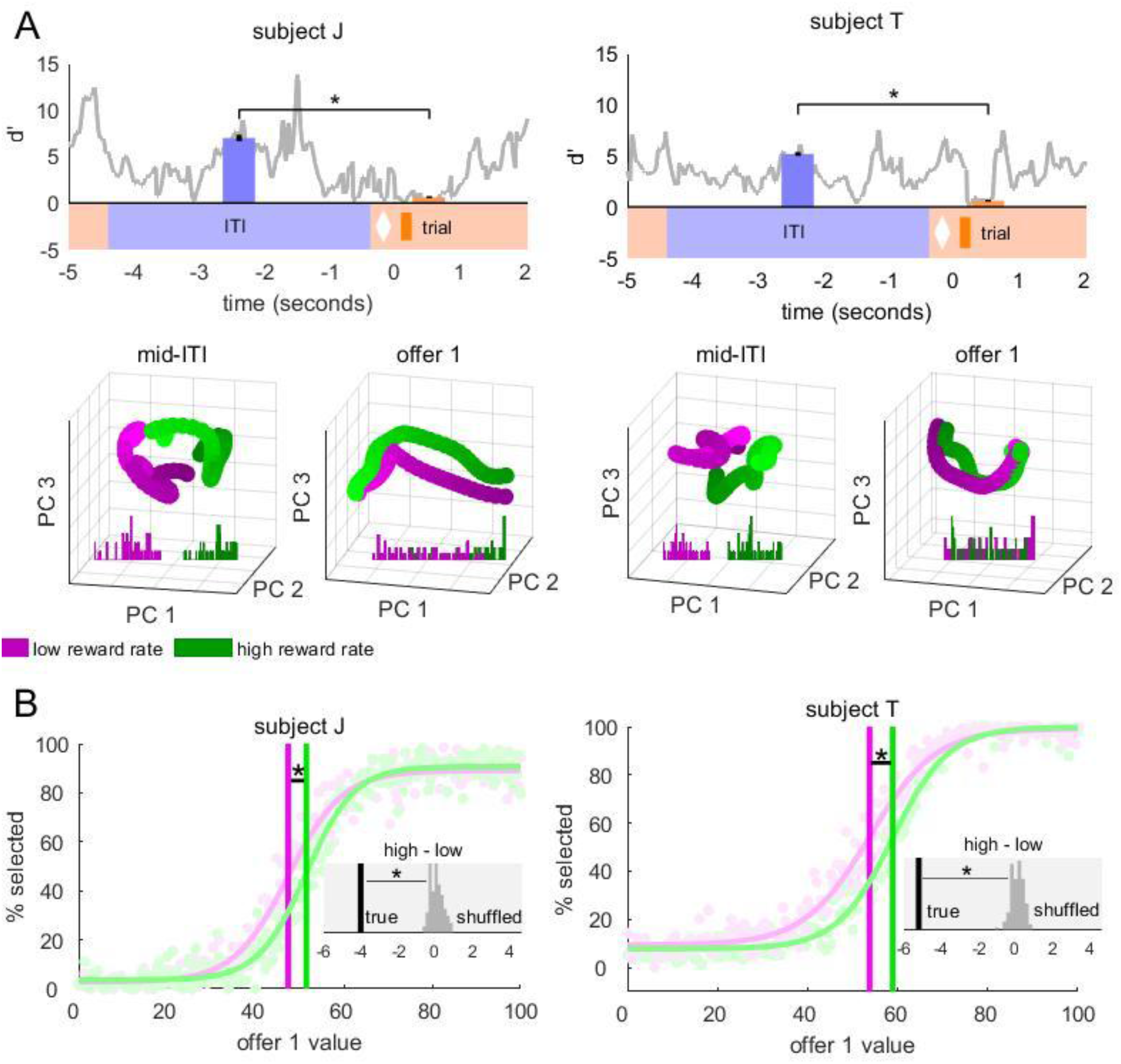
Discriminability of reward rate is higher at ITI than task-on period, and subjects behaviorally exploit reward rate information. **(A)** Upper panels: Discriminability of high versus low reward rate in top 3 PCs over time. Discriminability (d’) at the midpoint of the ITI epoch is significantly higher than d’ at the midpoint of the trial epoch in both subjects. Error bars represent standard deviation. Lower panels: Projection of principal components for high (green) and low (magenta) reward rates into a 3D space, where each point represents the top three principal components resulting from a PCA conducted on a 500 ms period centered around one time point. There are 50 time points for each condition, describing the 500 ms epoch centered on the midpoint of the ITI and trial epochs. Histograms depict the distances of each point from the plane that separates the two groups of points, as determined by linear discriminant analysis. **(B)** Subjects’ behavioral threshold for accepting an offer changes as recent reward rate changes: behavior threshold is higher for high reward rates than low reward rates for both subjects. Insets: the difference between the high reward rate threshold and low reward rate thresholds is significantly greater than the difference resulting from shuffled data.

We were interested in the question of whether information is more separable (i.e., decodable) during the ITI than it is during the trial epoch (Cohen & Maunsell, 2010). To determine the separability between high-reward-rate and low-reward-rate trials, we used a linear discriminant analysis (LDA) and calculated the distance (*d’*) between the cloud of high-reward-rate points and that of low-reward-rate points. A larger distance means a greater difference in the way the vmPFC population represents high vs. low reward rate. In other words, a downstream brain region with access to this information would have an easier time determining whether reward rate was high or low with a high *d’*. We found that high and low reward rate trials were significantly more separable during the ITI than during the trial in both subjects (subject J: ITI epoch *d’* 6.97, trial epoch *d’* 0.63; p<0.001, paired t-test; subject T: ITI epoch *d’* 5.17, trial epoch *d’* 0.61, p < 0.001, paired t-test; **Figure 2A, Methods**).

If the global reward rate is actively calculated during the ITI, then it should influence choice behavior during foraging. We previously reported that the behavioral threshold does change systematically with reward rate (Mehta et al., 2019). Here we demonstrate a significant difference between behavioral threshold over the same categories we found separation between during the ITI: high and low reward rate (subject J: low reward rate threshold 47.41%, high reward rate threshold 51.50%, p< 0.01, 100-repetition bootstrap; subject T: low reward rate threshold 53.73%, high reward rate threshold 58.89%; p<0.01,100-repetition bootstrap; **Figure 2B, Methods**). This illustrates a potential use for distinguishing high and low reward rate before the trial starts: adjusting behavioral strategy in accordance with reward rate.

### Dimensionality of neural subspace is greatest during the inter-trial-interval

We next examined the relationship between ITI and trial epoch ensemble ***dimensionality***. Ensembles of neurons have constrained firing rate patterns; that is, single neurons have a limited range of firing rates they can take and this range is determined in part by the responses of other neurons. That range, collectively, defines the neural manifold or subspace of the population (Gallego et al., 2017; Fusi et al., 2016). Dimensionality, then, refers to the number of the axes within the larger space spanned by the subspace. It is a useful measure because it provides a potential proxy for the complexity of processing going on within an area. Among other things, greater dimensionality also allows for more ready separation of response patterns, thus allowing for more abstract processing and learning (Fusi et al., 2016).

There are many ways to quantify the dimensionality of a subspace. We used a previously developed method (Machens et al., 2010; Rouse & Schieber, 2018). Specifically, we defined dimensionality as the number of principal components it takes to explain 90% of the variance in the neural activity at any particular moment (**Figure 3A**). We found that dimensionality is substantially greater during the ITI epoch than during the trial epoch in both subjects (subject J: 10.43 dimensions to 5.11 dimensions; p< 0.001, paired-t-test; subject T: 10.90 dimensions to 5.32 dimensions, p<0.001, paired t-test). (**Figure 3B**, upper panels). To compute these values and test significance, we conducted a bootstrap in which we left out 25% of our sample each time we computed this dimensionality value (using the remaining 75%) and averaged these values (**Figure 3B**, lower panels).

**Figure 3.**
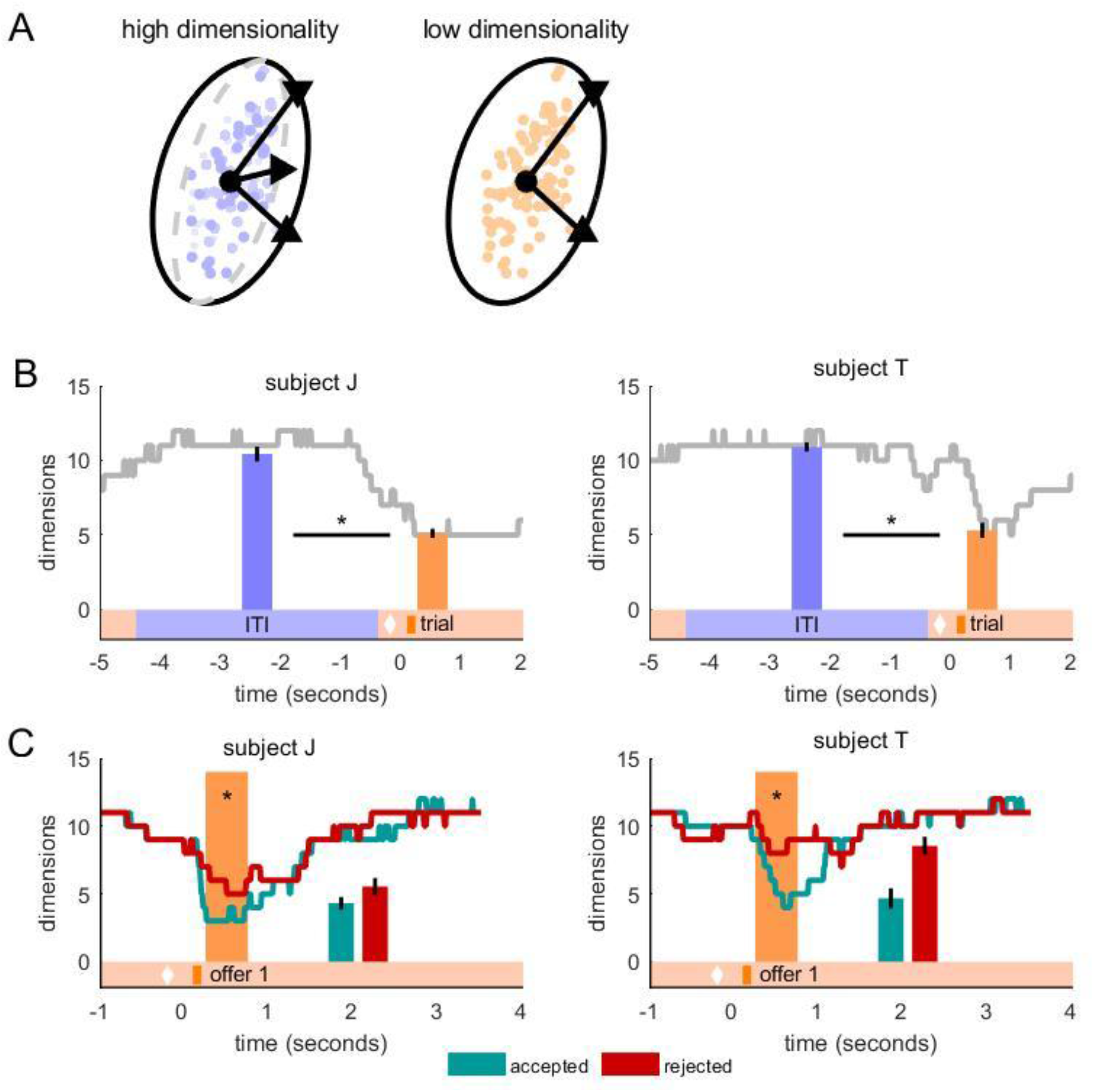
Subspace dimensionality is higher in the ITI than during the trial. **(A)** Schematic representing high and low dimensionality subspaces. **(B)** Dimensionality is significantly higher during the ITI epoch than the trial epoch for both subjects. Each point on gray line represents dimensionality of subspace over the preceding 500 ms. Blue and orange bars represent the dimensionality values centered on the ITI epoch and trial epoch generated by bootstrapped data. Error bars are standard deviation. **(C)** Dimensionality is significantly lower for accepted offers (teal) than rejected offers (red) for both subjects. Teal and red lines represent dimensionality at each point in time as in (B). Bars represent dimensionality centered on the midpoint of the trial epoch for accepted and rejected offers, using bootstrapped data. Error bars are standard deviation.

To gain deeper insight into these processes, we next asked whether this change in dimensionality is limited to the ITI vs. task periods. We hypothesized that if it reflects variations in focus, we should be able to see changes as focus changes within the task itself (cf. Hayden et al., 2009). For example, post-reward periods should have greater task focus than search periods because they involve greater local focus (on consumption), while search periods involve relatively more global focus (because they involve a search-like contemplation of multiple options). Our data are consistent with this hypothesis. We compared the dimensionality of the trial epoch between accepted and rejected offers using the same method specified in Figure 3B. Viewing an accepted offer (which led to a reward consumption period) resulted in a significantly lower dimensionality than viewing a rejected offer (which continued the trial); the effect was observed in both subjects (**Figure 3C**, subject J: accepted offers 4.31 dimensions, rejected offers 5.54 dimensions, p < 0.001, unpaired T-test; subject T: accepted offers 4.67 dimensions, rejected offers 8.54 dimensions, p<0.001, unpaired T-test).

### Reduction in dimensionality is not explained by task-driven firing rate changes

The observed reduction in dimensionality could potentially be explained by some other factor, such as the increase in firing at specific phases of the trial across all neurons. For example, a few neurons dramatically increasing their firing rate at stimulus onset could compress the subspace dimensionality, even though the effect would not be reflective of processing. Indeed, we found that average firing rate significantly increases between the ITI and task (significantly in subject J: mid-ITI normalized mean firing rate -1.03 spikes/second, offer 1 firing rate 1.39 spikes/second, p<0.001, paired t-test; subject T: mid-ITI normalized mean firing rate - 0.09, offer 1 normalized firing rate 0.00, p =0.01, paired t-test, data smoothed over 20 bins;

To test for the possibility that firing rate changes are responsible for the dimensionality changes we see, we created control data that had the same mean firing rate as the real data, but scrambled relationships between individual neurons **(methods)**. In both subjects, the shuffled dimensionality is significantly higher than the true dimensionality (subject J: 7.57 dimensions versus 5.35 dimensions, p<0.001; subject T: 8.30 vs 7.33 dimensions, p, = 0.003). This result suggests that while some of the reduction in dimensionality could be due to the trial-driven firing rate transient, a large part of it is independent of that.

### Reorganization rate of neural subspace reorganization is greatest during the inter-trial-interval

The subspace an ensemble response occupies can change. This change, which can implement functional partition, is often an indicator of changing function roles of the population (Elsayed et al., 2016; Jiang et al., 2020; Tang et al., 2020; Yoo & Hayden, 2020) Here, we refer to the amount of subspace reorganization at each point in time as subspace *reorganization rate* - where a high amount of reorganization from one moment to the next reflects high change rate (and its converse, low stability). Changes in neural activity patterns during rest states are generally considered to be slow (<0.1 Hz; Cabral et al., 2014; Deco & Jirsa, 2012; de Pasquale & Marzetti, 2019). That is, techniques ranging from functional magnetic resonance imaging to magnetoencephalography, to electrophysiology, have revealed that low-frequency patterns characterize neural activity during rest. As a result, one would expect the pattern of changes we find during the ITI to be slow as well. However, there is emerging evidence of rapid computations during rest broadly across the default mode network (Baker et al., 2014; Khanna et al., 2015), and specifically in the motor cortex (Vidaurre et al., 2016). Here we provide evidence of fast-changing neural patterns (significant reorganization of the neural subspace between 500 ms epochs) during rest in the ventromedial prefrontal cortex. These results point to a changing view of default mode activity as not only slow oscillatory activity patterns, but also fast-changing dynamic activity.

To assess the subspace reorganization rate, we measured the similarity of the eigenvectors between each pair of subspaces before and after every given time point (10 ms intervals) of the task with 500 ms epochs (**Methods**). We considered every timepoint from the beginning of the ITI through two seconds after the reveal of the first offer. For each timepoint, we defined two time periods - 500 ms before, and 500 ms after - and computed the PCA coefficients for both time periods. We then projected the coefficients from the *before* time period into the *after* subspace and determined the percentage of *after* variance that was explained by the *before* coefficients (Elsayed et al., 2016; Yoo & Hayden, 2020; Jiang et al., 2020; Khanna et al., 2019). This overlap, expressed as a percentage, indicates the amount of reorganization of the neural subspace. A reorganization rate of 100% means the *before* subspace coefficients explained none of the *after* subspace (fast reorganization), and thus has completely reorganized, while 0% velocity means the *before* subspace explains all of the *after* subspace, and thus has not reorganized at all.

We found that the subspace reorganization rate is significantly greater during the ITI than during the task for both subjects (**Figure 4B**). Specifically, in subject J the reorganization for sequential epochs at the midpoint of the ITI was 66.59% but only 28.34% at the midpoint of the trial epoch (p<0.001, one-sample t-test between true difference and shuffled differences, see **Methods**). For subject T, these numbers were 72.76% for the ITI epoch and 55.35% for the trial epoch (p<0.001, one-sample t-test between true difference and shuffled differences). In other words, vmPFC neuronal subspace occupancy changes much more over time during the ITI than during the task; once the trial begins, however, the neural subspace becomes much more stable.

**Figure 4.**
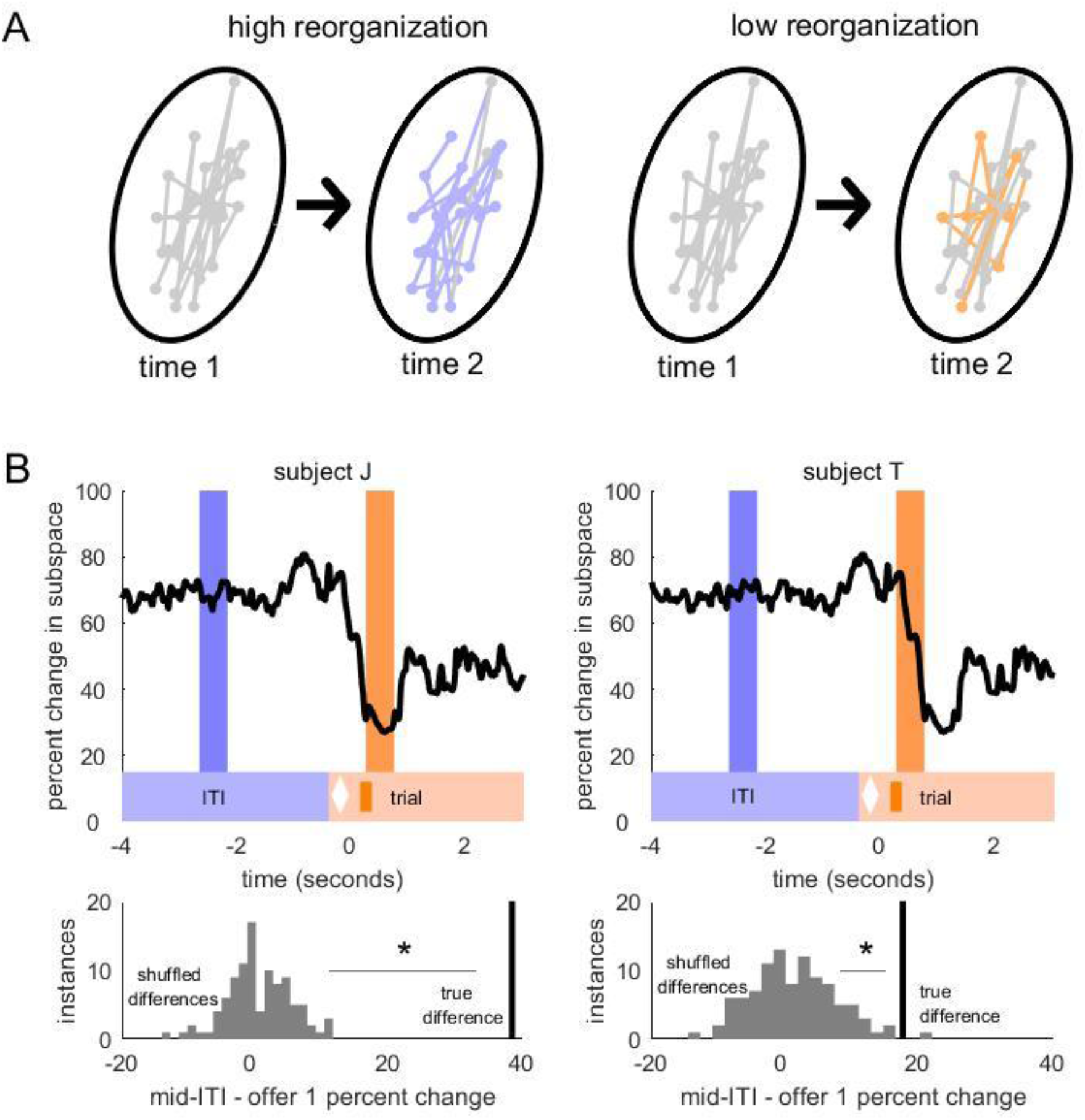
Subspace reorganizes much more during ITI than during trial. **(A)** Schematic of high subspace reorganization versus low subspace reorganization. **(B)** Upper panels: The percent change in subspace over the midpoint of the ITI epoch is greater than that over the midpoint of the trial epoch in both subjects. Lower panels: this difference is significantly greater than the distribution of differences generated by shuffling data 100 times.

Finally, we note that the ITI subspace is organized drastically differently from the trial subspace. That is, we conducted the same projection analysis on the 500 ms epoch centered around the midpoint of the ITI and the 500 ms epoch centered around the midpoint of the trial epoch, and found that the PC space of the ITI epoch changes 83.07% in subject J and 76.76% in subject T.

## DISCUSSION

The ITI, and globally focused cognition more generally, occupies a large amount of our lives and yet has not been studied nearly as much as task-oriented locally focused cognition. Indeed, despite its quotidian nature, it remains strangely inaccessible to many conventional cognitive neuroscientific methods (Raichle, 2006). Nonetheless, studies of default mode processing have amply demonstrated that the time between trials and the times of global cognition more generally are not simply “idling” or “paused” time in the brain (Boly et al., 2007; Konishi et al., 2015; Weissman et al., 2006; Smallwood et al., 2013;, Baird et al., 2013). Instead, the brain makes use of this time to engage in a variety of cognitive functions. Many of these seem to be related to mnemonic processing.

Here, we examined the responses of a population of neurons in area 14 of the vmPFC while macaques performed a foraging search task (Mehta et al., 2019). We imposed an unusually long interval between sequential trials (duration of 4 seconds), a design feature that allowed us to compare neuronal ensemble responses during task-on (trial) and task-off (inter-trial interval, ITI) periods while avoiding immediate post-trial and pre-trial effects. We examined encoding of a key piece of mnemonic information in the task - recent reward rate, which we show are attended and learned. During the ITI, that information is encoded in a more separable format: an effect we demonstrate with our discriminability analysis. In other words, information is present during both the task and ITI epochs, but it is more untangled during the ITI (DiCarlo et al., 2012; Yoo & Hayden, 2018; DiCarlo & Cox, 2007). This finding suggests that the brain actively reformats task-relevant information depending on task context and, in particular, that it makes mnemonic information more functionally available during the ITI than during the task itself.

Why would reward rate information be more separable during the ITI than during the task itself? During the trial, the subject must do two things - (1) apply learned information for purposes of guiding decisions and (2) be alert for new incoming information. Both processes may be antagonistic to the kinds of neural processes that are associated with mnemonic processing, including computing and representing the recent average reward value. In other words, explicit reward value representation is a guide for behavior but it is conceptually quite distinct from its implementation, and cognition may have a finite capacity. It may be efficient, then, to process it asynchronously during low-demand periods. Indeed, we hypothesize that the more accessible format during the delay reflects the need to process it and develop a decision strategy; that strategy only needs to be implemented during choice, so the information that guides the strategy can be stored in a more latent manner.

We also found that the ITI is associated with more complex processing, as indicated by two complementary measures: subspace dimensionality and subspace change rate. Dimensionality refers to the volume occupied by an ensemble in a high-dimensional space: a higher dimension is presumed to reflect less redundancy and more complex and multifarious response patterns (Cunningham & Byron, 2014). Subspace change rate, on the other hand is a new measure that we introduce here, but it is one that follows naturally from past work on subspace reorganization - it is, essentially the rate at which reorganization occurs. We reasoned that faster reorganization may imply rapid cognitive processing with multiple changing priorities. Some of the dimensionality and velocity results are likely related to the specific mnemonic function discussed above, but some likely reflect even more complex processing to which we don’t have experimental access.

Why would the brain alternate between processing different types of information in different modes? We speculate that the brain is not a strictly parallel processor, and that its capacity is highly restricted (Shenav et al., 2017). Specifically, we conjecture that broad and narrow-focused attention naturally competes and cannot be effectively implemented at the same time. In other words, we propose that information processing is analogous to preemptive multiple processing in computer science. This type of multiple processing operates by prioritizing the task and context switch operation allows the interruption of certain tasks and resumes later. The global information processing can be interrupted by task engagement (local phase) and resumed once the individual occasion of search ends. According to preemptive multiple processing, the operating processor never shuts down but they switch the information being processed.

## METHODS

We conducted all procedures in compliance with the Public Health Service’s Guide for the Care and Use of Animals and approved by the University Committee on Animal Resources at the University of Rochester. These data were previously analyzed in a published study (Mehta et al., 2019). Subjects were two male rhesus macaques (*Macaca mulatta*: subject J age 10 years; subject T age 5 years). To maintain head position, we used a small titanium prosthesis. We trained the subjects first to habituate to laboratory conditions and then to perform oculomotor tasks for liquid reward. To record neural data, we placed a Cilux recording chamber (Crist instruments) over the ventromedial prefrontal cortex (vmPFC) of each animal. We verified the accuracy of this position using magnetic resonance and a Brainsight system (Rogue Research Inc.). We provided the animals with all appropriate analgesics and antibiotics post-procedure, and regularly sterilely cleaned and sealed the implanted chamber.

### Recording site

We performed all recordings between morning and midday (10am to 3pm), using a standard recording grid (Crist Instruments) to approach area 14 (vmPFC) as described by the Paxinos atlas (Paxinos et al., 2000). Specifically, we recorded from neurons located between 42 and 31 mm rostral to the interaural plane, 0 and 7 mm horizontally from the brain’s ventral surface, and 0 to 7 mm sagittal from the medial wall.

### Electrophysiological techniques, eye tracking and reward delivery

All methods used have been described previously and are summarized here (Strait et al., 2014). We used a microdrive (NAN instruments) to guide single electrodes (Frederick Haer & Co., impedance range 0.7 to 5.5 MU). Each recording session, we isolated between one and four neurons on a Plexon system (Plexon, Inc.). We used data from all neurons we could isolate for at least 300 consecutive trials, regardless of their behavior during the task. In practice, 86% of neurons had over 500 trials

To obtain eye position data at 1,000 Hz, we used an infrared eye-monitoring camera system (SR Research). We controlled the task presentation with a computer running Matlab (Mathworks) with Psychtoolbox and Eyelink Toolbox. Our visual stimuli were colored shapes on a computer monitor located 60 cm from the animal at eye level. We controlled liquid reward delivery with a standard solenoid valve.

### Experimental Design

Subjects performed a diet-search task (Figure 1) that is described in detail in a previous publication (Mehta et al., 2019). The task is a conceptual extension of previous foraging tasks we have developed. Each trial, subjects were presented with either four or seven white diamonds that appeared in random positions on the screen. 400 ms of continuous fixation on one of these diamonds caused it to disappear and reveal a reward offer underneath. Reward offers were partially-filled-in orange bars that indicated riskless reward amount by the proportion of their area that was filled in. There were 200 possible reward offer sizes per offer. To accept an offer, subjects had to continuously fixate on one for 300 ms.

Subjects freely searched through the diamonds in any order, but had to accept an offer to obtain a reward and end the trial. Rejection of an offer (breaking fixation before 300 ms had elapsed) resulted in a continuation of the trial (free search through the diamonds). Reward was delivered immediately after acceptance of an offer, which was followed by a four-second inter-trial-interval.

Previous training history for these subjects included two types of foraging tasks (Blanchard and Hayden, 2015; Blanchard et al., 2015), complex choice tasks with time elements (Blanchard et al., 2014), two types of gambling tasks (Farashahi et al., 2018; Azab and Hayden, 2017), attentional tasks (similar to those in Hayden and Gallant, 2013), and two types of reward-based decision tasks (Pirrone et al., 2018; Wang and Hayden, 2017).

### Statistical Analysis for Physiology

We formed Peri-Stimulus Time Histograms (PSTHs) by aligning spike rasters to each offer reveal of each trial and averaging firing rates across multiple trials. Firing rates were normalized (where indicated) by subtracting the mean and dividing by the standard deviation of the entire neuron’s PSTH.

### Analysis epochs

Before beginning analysis, we determined two epochs over the course of the trial to correspond to the ITI and the task. We selected the 500 ms period in the middle of the 4-second ITI to represent the *ITI epoch*, and the 500ms in the middle of the 1-second epoch following the reveal of the first offer to represent the *offer epoch*.

### Creating the PC spaces

To create a PC-space for the ITI and first offer of each trial, we began with the PSTHs of all the trials, sorted into two groups: trials where the subject accepted the first offer, and trials where the subject rejected the first offer. These PSTHs were Gaussian smoothed over 15 time bins (150 ms), and normalized. (Note that while data was divided into accept and reject data for the dimensionality and orthogonality analyses, it was divided into low and high reward rate datasets for the discriminability analysis).

We then found the mean PSTH for each cell for each of these groups, giving us two matrices per subject - each with cells as rows and time points of the PSTH as columns. In other words, we computed the average behavior of each cell for trials where the subject accepted the first offer, and trials where the subject rejected the first offer. We then appended the *reject* matrix to the *accept* matrix to create a combined matrix.

To obtain PC spaces over the course of the ITI and first offer, we used a sliding boxcar of 50 time bins (500 ms) to perform a PCA on each 500-ms epoch of PSTH data from the beginning of the ITI through 2 seconds following the reveal of the first offer.

To obtain PC spaces for trials following the acceptance or rejection of an offer, we used a sliding boxcar of 50 time bins to perform a PCA on each 500-ms epoch of PSTH data from 2 seconds before the reveal of each offer to 3 seconds after.

### Discriminability analysis

We defined *discriminability* between two task conditions as the separation between representations of those variables in principal component space. To this end, we performed PCAs as described above, using neural data with each trial split into either a low-reward rate (between 0 and 71% for subject J; 0 and 72% for subject T, see Mehta 2019 for calculation of reward rate), or high-reward-rate (71%-100% for subject J; 72% and above for subject T) condition. We then obtained the top three principal components (the principal components that explain the three largest amounts of variance in the neural activity) for each condition and used a linear discriminant analysis (LDA) to assess their separability.

Specifically, we plotted the top three principal components for each condition over a specific time epoch, so that we obtained a cloud of three-dimensional points representing all the time points in one epoch for each condition. Then, the LDA provided us with a plane separating the two clouds of points. To obtain *d’*, the separability between these two clouds, we first calculated the distance of each of these points from the separating plane, as well as the standard deviations of the distributions of distances for each condition. We defined *d’* as follows:

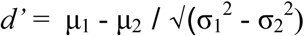

Where µ_1_ and µ_2_ are the means of the distances for each condition (low and high reward rate) and σ_1_ are σ_2_ are the standard deviations of these distributions.

### Dimensionality analysis

We defined *dimensionality* of an epoch as the number of principal components it takes to explain 90% of the variance in the neural activity during that epoch (Elsayed et al., 2016; Yoo &Hayden, 2020). We used the PCAs run as detailed above on our epochs of interest and obtained the percentage of the variance that each PC explained, sorted from greatest variance explained to least. For each epoch of interest, we found the number of principal components, starting with the one that explained the most variance, that it took to explain a minimum of 90% of the total variance. This value we termed the ‘dimensionality.’

It is important to note that dimensionality can only be a whole number: it represents a number of principal components. When we report dimensionality as having decimal values, it is because we have averaged several whole numbers over a period of time. We interpret these decimal values as estimates of dimensionality.

To perform significance tests on dimensionality data, we had to bear in mind that each sequential time point’s dimensionality is created from data that overlaps with the next time point. Thus, we could not perform a simple t-test between, for example, the dimensionalities of the time points between the reveal of the first offer and one second following. Instead, we used a bootstrap: we selected the central time point of each analysis epoch (which represents data from 250 ms before that point and 250 ms after) and recalculated it 100 times using 75% of the original data each time. We then performed a t-test between these two groups of 100 points between epochs.

To create control data, we randomly swapped the PSTHs of the *offer 1* epoch between cells. For example, if there were three real cells, Cell A, B and C and three control cells, Cell A’, B’, and C, Cell A’ may have the *offer 1* epoch of Cell C’, Cell B’ might have the *offer 1* epoch of Cell A, and Cell C would have the *offer 1* epoch of Cell B. Thus, the control data would have the same mean firing rate as the real data. However, suppose that the *combination* of Cells A and B specifically is meaningful to the vmPFC; that is, population information is meaningful. Now, the combination of Cell A and Cell B’s firing rate is meaningless, because their PSTHs have been swapped out with those of other cells. Thus, our control data was such that the mean firing rate at every time point of the *offer 1* epoch was the same, but the properties of the data at the population level are meaningless.

### Orthogonality analysis

To estimate the amount of reorganization between subspaces for the two epochs, we did the following (Elsayed et al., 2016; Yoo & Hayden, 2020). We first performed principal components analysis (PCA) on the matrix P1 for each epoch (see above) to obtain the epoch-1 PCs. For the remainder of the analysis, we used only the top ten PCs resulting from each PCA. We obtained the top ten epoch 2 PCs by performing PCA on matrix P2. To examine the relationship between the subspaces, we projected the epoch 1 activity onto the epoch 2 PCs and quantified the percent of variance explained relative to the total variance of epoch 1.

To perform significance tests on orthogonality data, we noted that orthogonality values are derived from principal components that describe overlapping epochs of neural data. Thus again, we could not perform a simple t-test between the average orthogonality of different epochs. Instead, we created shuffled data by scrambling the PCA inputs between the two epochs of interest, 100 times. We then calculated the orthogonality of the central time point of the ITI and the offer epoch 100 times with these sets of scramble data. We compared the true difference in orthogonality to the distribution of shuffled differences with a t-test to obtain a p-value.

## Acknowledgements

This work is supported by a R01 from NIH (DA037229) to BYH. We thank Meghan Castagno, Marc Mancarella, and Giuliana LoConte for assistance with data collection, and the rest of the Hayden lab for valuable discussions.

## REFERENCES

Azab, H., & Hayden, B. Y. (2017). Correlates of decisional dynamics in the dorsal anterior cingulate cortex. PLoS biology, 15(11), e2003091.

Baird, B., Smallwood, J., Gorgolewski, K. J., & Margulies, D. S. (2013). Medial and lateral networks in anterior prefrontal cortex support metacognitive ability for memory and perception. Journal of Neuroscience, 33(42), 16657–16665.

Baker, A. P., Brookes, M. J., Rezek, I. A., Smith, S. M., Behrens, T., Smith, P. J. P., & Woolrich, M. (2014). Fast transient networks in spontaneous human brain activity. Elife, 3, e01867.

Behrens, T. E., Muller, T. H., Whittington, J. C., Mark, S., Baram, A. B., Stachenfeld, K. L., & Kurth-Nelson, Z. (2018). What is a cognitive map? Organizing knowledge for flexible behavior. Neuron, 100(2), 490–509.

Blanchard, T. C., Strait, C. E., & Hayden, B. Y. (2015). Ramping ensemble activity in dorsal anterior cingulate neurons during persistent commitment to a decision. Journal of neurophysiology, 114(4), 2439–2449.

Blanchard, T. C., Hayden, B. Y., & Bromberg-Martin, E. S. (2015). Orbitofrontal cortex uses distinct codes for different choice attributes in decisions motivated by curiosity. Neuron, 85(3), 602–614.

Blanchard, T. C., & Hayden, B. Y. (2015). Monkeys are more patient in a foraging task than in a standard intertemporal choice task. PloS one, 10(2), e0117057.

Boly, M., Balteau, E., Schnakers, C., Degueldre, C., Moonen, G., Luxen, A., … & Laureys, S. (2007). Baseline brain activity fluctuations predict somatosensory perception in humans. Proceedings of the National Academy of Sciences, 104(29), 12187–12192.

Cabral, J., Kringelbach, M. L., & Deco, G. (2014). Exploring the network dynamics underlying brain activity during rest. Progress in neurobiology, 114, 102–131.

Cohen, M. R., & Maunsell, J. H. (2010). A neuronal population measure of attention predicts behavioral performance on individual trials. Journal of Neuroscience, 30(45), 15241–15253.

Cole, M. W., Ito, T., Bassett, D. S., & Schultz, D. H. (2016). Activity flow over resting-state networks shapes cognitive task activations. Nature neuroscience, 19(12), 1718–1726.

Cunningham, J. P., & Byron, M. Y. (2014). Dimensionality reduction for large-scale neural recordings. Nature neuroscience, 17(11), 1500–1509.

Daselaar, S. M., Prince, S. E., & Cabeza, R. (2004). When less means more: deactivations during encoding that predict subsequent memory. Neuroimage, 23(3), 921–927.

Deco, G., & Jirsa, V. K. (2012). Ongoing cortical activity at rest: criticality, multistability, and ghost attractors. Journal of Neuroscience, 32(10), 3366–3375.

de Pasquale, F., & Marzetti, L. (2019). Temporal and spectral signatures of the default mode network. Magnetoencephalography: From Signals to Dynamic Cortical Networks, 571–603.

DiCarlo, J. J., & Cox, D. D. (2007). Untangling invariant object recognition. Trends in cognitive sciences, 11(8), 333–341.

DiCarlo, J. J., Zoccolan, D., & Rust, N. C. (2012). How does the brain solve visual object recognition?. Neuron, 73(3), 415–434.

Elsayed, G. F., Lara, A. H., Kaufman, M. T., Churchland, M. M., & Cunningham, J. P. (2016). Reorganization between preparatory and movement population responses in motor cortex. Nature communications, 7(1), 1–15.

Farashahi, S., Azab, H., Hayden, B., & Soltani, A. (2018). On the flexibility of basic risk attitudes in monkeys. Journal of Neuroscience, 38(18), 4383–4398.

Ferguson, T. S. (1989). Who solved the secretary problem?. Statistical science, 4(3), 282–289.

Fusi, S., Miller, E. K., & Rigotti, M. (2016). Why neurons mix: high dimensionality for higher cognition. Current opinion in neurobiology, 37, 66–74.

Gallego, J. A., Perich, M. G., Miller, L. E., & Solla, S. A. (2017). Neural manifolds for the control of movement. Neuron, 94(5), 978–984.

Hayden, B. Y., Nair, A. C., McCoy, A. N., & Platt, M. L. (2008). Posterior cingulate cortex mediates outcome-contingent allocation of behavior. Neuron, 60(1), 19–25.

Hayden, B. Y., Pearson, J. M., & Platt, M. L. (2009). Fictive reward signals in the anterior cingulate cortex. science, 324(5929), 948–950.

Hayden, B. Y., Smith, D. V., & Platt, M. L. (2009). Electrophysiological correlates of default-mode processing in macaque posterior cingulate cortex. Proceedings of the National Academy of Sciences, 106(14), 5948–5953.

Hayden, B. Y., Pearson, J. M., & Platt, M. L. (2011). Neuronal basis of sequential foraging decisions in a patchy environment. Nature neuroscience, 14(7), 933.

Hayden, B., & Gallant, J. (2013). Working memory and decision processes in visual area v4. Frontiers in neuroscience, 7, 18.

Hayden, B. Y., & Platt, M. L. (2010). Neurons in anterior cingulate cortex multiplex information about reward and action. Journal of Neuroscience, 30(9), 3339–3346.

Heilbronner, S. R., & Hayden, B. Y. (2016). Dorsal anterior cingulate cortex: a bottom-up view. Annual review of neuroscience, 39, 149–170.

Heilbronner, S. R., & Hayden, B. Y. (2016). The description-experience gap in risky choice in nonhuman primates. Psychonomic bulletin & review, 23(2), 593–600.

Heilbronner, S. R. (2017). Modeling risky decision-making in nonhuman animals: shared core features. Current opinion in behavioral sciences, 16, 23–29.

Jiang, X., Saggar, H., Ryu, S. I., Shenoy, K. V., & Kao, J. C. (2020). Structure in Neural Activity during Observed and Executed Movements Is Shared at the Neural Population Level, Not in Single Neurons. Cell reports, 32(6), 108006.

Jilka, S. R., Scott, G., Ham, T., Pickering, A., Bonnelle, V., Braga, R. M., … & Sharp, D. J. (2014). Damage to the salience network and interactions with the default mode network. Journal of neuroscience, 34(33), 10798–10807.

Kao, T. C., Sadabadi, M. S., & Hennequin, G. (2020). Anticipatory control of movement in a thalamo-cortical circuit model. bioRxiv.

Keemink, S. W., & Machens, C. K. (2019). Decoding and encoding (de) mixed population responses. Current Opinion in Neurobiology, 58, 112–121.

Khanna, A., Pascual-Leone, A., Michel, C. M., & Farzan, F. (2015). Microstates in resting-state EEG: current status and future directions. Neuroscience & Biobehavioral Reviews, 49, 105–113.

Khanna, S. B., Snyder, A. C., & Smith, M. A. (2019). Distinct sources of variability affect eye movement preparation. Journal of Neuroscience, 39(23), 4511–4526.

Konishi, Mahiko, Donald George McLaren, Haakon Engen, and Jonathan Smallwood. “Shaped by the past: the default mode network supports cognition that is independent of immediate perceptual input.” PloS one 10, no. 6 (2015): e0132209.

Machens, C. K., Romo, R., & Brody, C. D. (2010). Functional, but not anatomical, separation of “what” and “when” in the prefrontal cortex. Journal of Neuroscience, 30(1), 350–360.

Mason, M. F., Norton, M. I., Van Horn, J. D., Wegner, D. M., Grafton, S. T., & Macrae, C. N. (2007). Wandering minds: the default network and stimulus-independent thought. Science, 315(5810), 393–395.

Mazoyer, B., Zago, L., Mellet, E., Bricogne, S., Etard, O., Houdé, O., … & Tzourio-Mazoyer, N. (2001). Cortical networks for working memory and executive functions sustain the conscious resting state in man. Brain research bulletin, 54(3), 287–298.

Mehta, P. S., Tu, J. C., LoConte, G. A., Pesce, M. C., & Hayden, B. Y. (2019). Ventromedial prefrontal cortex tracks multiple environmental variables during search. Journal of Neuroscience, 39(27), 5336–5350.

Northoff, G., Qin, P., & Nakao, T. (2010). Rest-stimulus interaction in the brain: a review. Trends in neurosciences, 33(6), 277–284.

Paxinos, G., Huang, X. F., & Toga, A. W. (2000). The rhesus monkey brain in stereotaxic coordinates.

Pirrone, A., Azab, H., Hayden, B. Y., Stafford, T., & Marshall, J. A. (2018). Evidence for the speed–value trade-off: Human and monkey decision making is magnitude sensitive. Decision, 5(2), 129.

Raichle, M. E. (2006). The brain’s dark energy. Science-New York Then Washington-, 314(5803), 1249.

Raichle, M. E. (2010). Two views of brain function. Trends in cognitive sciences, 14(4), 180–190.

Raichle, M. E. (2015). The brain’s default mode network. Annual review of neuroscience, 38, 433–447.

Rouse, A. G., & Schieber, M. H. (2018). Condition-dependent neural dimensions progressively shift during reach to grasp. Cell reports, 25(11), 3158–3168.

Shenhav, A., Musslick, S., Lieder, F., Kool, W., Griffiths, T. L., Cohen, J. D., & Botvinick, M. M. (2017). Toward a rational and mechanistic account of mental effort. Annual review of neuroscience, 40, 99–124.

Smallwood, J., Tipper, C., Brown, K., Baird, B., Engen, H., Michaels, J. R., … & Schooler, J. W. (2013). Escaping the here and now: evidence for a role of the default mode network in perceptually decoupled thought. Neuroimage, 69, 120–125.

Stephens, D. W., & Krebs, J. R. (1986). Foraging theory (Vol. 1). Princeton University Press.

Strait, C. E., Blanchard, T. C., & Hayden, B. Y. (2014). Reward value comparison via mutual inhibition in ventromedial prefrontal cortex. Neuron, 82(6), 1357–1366.

Sugrue, L. P., Corrado, G. S., & Newsome, W. T. (2004). Matching behavior and the representation of value in the parietal cortex. science, 304(5678), 1782–1787.

Tang, C., Herikstad, R., Parthasarathy, A., Libedinsky, C., & Yen, S. C. (2020). Minimally dependent activity subspaces for working memory and motor preparation in the lateral prefrontal cortex. eLife, 9, e58154.

Wang, M. Z., & Hayden, B. Y. (2017). Reactivation of associative structure specific outcome responses during prospective evaluation in reward-based choices. Nature communications, 8(1), 1–13.

Weissman, D. H., Roberts, K. C., Visscher, K. M., & Woldorff, M. G. (2006). The neural bases of momentary lapses in attention. Nature neuroscience, 9(7), 971–978.

Yoo, S. B. M., & Hayden, B. Y. (2018). Economic choice as an untangling of options into actions. Neuron, 99(3), 434–447.

Yoo, S. B. M., & Hayden, B. Y. (2020). The transition from evaluation to selection involves neural subspace reorganization in core reward regions. Neuron, 105(4), 712–724.

Zou, Q., Ross, T. J., Gu, H., Geng, X., Zuo, X. N., Hong, L. E., … & Yang, Y. (2013). Intrinsic resting□state activity predicts working memory brain activation and behavioral performance. Human brain mapping, 34(12), 3204–3215.

